# Neutrophil extracellular trap stabilization by platelet factor 4 reduces thrombogenicity and endothelial cell injury

**DOI:** 10.1101/2023.01.09.522931

**Authors:** Anh T. P. Ngo, Amrita Sarkar, Irene Yarovoi, Nate D. Levine, Veronica Bochenek, Guohua Zhao, Lubica Rauova, M. Anna Kowalska, Kaitlyn Eckart, Nilam S. Mangalmurti, Ann Rux, Douglas B. Cines, Mortimer Poncz, Kandace Gollomp

**Affiliations:** Division of Hematology, Children’s Hospital of Philadelphia, Philadelphia, PA, USA; Department of Pediatrics, Perelman School of Medicine at the University of Pennsylvania, Philadelphia, PA, USA; Department of Medicine, Perelman School of Medicine at the University of Pennsylvania, Philadelphia, PA, USA; Department of Pathology and Laboratory Medicine, Perelman School of Medicine at the University of Pennsylvania, Philadelphia, PA, USA

**Keywords:** platelet factor 4, neutrophil extracellular trap, cell-free DNA, sepsis, thrombosis

## Abstract

Neutrophil extracellular traps (NETs) are abundant in sepsis, and proposed NET-directed therapies in sepsis prevent their formation or accelerate degradation. Yet NETs are important for microbial entrapment, as NET digestion liberates pathogens and NET degradation products (NDPs) that deleteriously promote thrombosis and endothelial cell injury. We proposed an alternative strategy of NET-stabilization with the chemokine, platelet factor 4 (PF4, CXCL4), which we have shown enhances NET-mediated microbial entrapment. We now show that NET compaction by PF4 reduces their thrombogenicity. In vitro, we quantified plasma thrombin and fibrin generation by intact or degraded NETs and cell-free (cf) DNA fragments, and found that digested NETs and short DNA fragments were more thrombogenic than intact NETs and high molecular weight genomic DNA, respectively. PF4 reduced the thrombogenicity of digested NETs and DNA by interfering, in part, with contact pathway activation. In endothelial cell culture studies, short DNA fragments promoted von Willebrand factor release and tissue factor expression via a toll-like receptor 9-dependent mechanism. PF4 blocked these effects. *Cxcl4^-/-^* mice infused with cfDNA exhibited higher plasma thrombin anti-thrombin (TAT) levels compared to wild-type controls. Following challenge with bacterial lipopolysaccharide, *Cxcl4^-/-^* mice had similar elevations in plasma TAT and cfDNA, effects prevented by PF4 infusion. Thus, NET-stabilization by PF4 prevents the release of short fragments of cfDNA, limiting the activation of the contact coagulation pathway and reducing endothelial injury. These results support our hypothesis that NET-stabilization reduces pathologic sequelae in sepsis, an observation of potential clinical benefit.

**HIGHLIGHTS:**

- In contrast to intact NETs, degraded NETs and cfDNA are prothrombotic and injure the endothelium.
- PF4 reduces the ability of degraded NETs and cfDNA to promote thrombosis and injure the endothelium.

## INTRODUCTION

Sepsis is a dysfunctional response to infection that leads to life-threatening organ damage. Although sepsis is a leading cause of mortality worldwide, treatment remains limited to antibiotics and supportive care.^1,2^ Over the past decade, multiple studies have shown that levels of plasma cell-free (cf) DNA are highly elevated in septic patient plasma and are associated with organ dysfunction and mortality.^3–6^ Recent studies have shown that cfDNA is not merely a marker of disease severity, but actively contributes to disease progression by activating the intrinsic pathway of coagulation and acting as a damage-associated molecular pattern (DAMP).^7^

Although a portion of cfDNA is derived from necrosis and apoptosis, in sepsis, the predominant source is neutrophil extracellular traps (NETs),^8^ webs of negatively-charged DNA complexed with positively-charged histones and antimicrobial proteins that capture and kill pathogens.^9–11^ When NETs are degraded by circulating nucleases (DNases), they release NET-degradation products (NDPs) including cfDNA, histones and neutrophil granule proteins that trigger prothrombotic pathways and cause oxidative vascular damage.^12–16^

Based on the finding that NDP levels correlate with end-organ damage and mortality in septic patients^3,5,17–20^ it has been proposed that preventing NETosis might be beneficial.^21,22^ However, NDP levels are markedly elevated in septic patients at clinical presentation,^17,23–25^ and blocking NET-release on admission may be ineffective. Another NET-based strategy proposed is to accelerate NET degradation by infusing DNases.^26–29^ However, studies of sepsis in murine models suggest that infusion of DNase I soon after infection liberates entrapped bacteria and increases levels of circulating NDPs, leading to worse outcomes.^30,31^

Platelet factor 4 (PF4, CXCL4) is a chemokine stored in platelet alpha granules and is 2-3% of total platelet releasate upon platelet activation. Local concentrations can exceed 12 μg/mL at sites of injury.^32^ Due to its strong positive charge, PF4 tetramers aggregate polyanions such as heparin^33^ and NETs.^13,31^ Previous work demonstrated that human (h) PF4 improves survival in the murine lipopolysaccharide (LPS) endotoxemia model of sepsis.^34^ We recently proposed and demonstrated that PF4 exerts this protective effect by binding to NETs, causing them to become physically compact and resist nuclease digestion.^31^ KKO, a monoclonal antibody directed against hPF4:polyanion complexes,^35^ binds PF4:NET complexes, further enhancing their resistance to nuclease digestion. We also showed that a deglycosylated version of KKO (DGKKO) works in concert with hPF4 to reduce NDP release and enhancing NET capture of bacteria, markedly improving survival in murine models of sepsis.^13,31,36^

It may seem counterintuitive to use hPF4 alone or with DGKKO to stabilize NETs in sepsis, as NETs are considered to be thrombogenic and injurious to the endothelial lining.^12–15,17^ However, the results of our in vitro studies are consistent with prior work that shows that unlike isolated DNA, intact NETs do not activate the contact pathway of coagulation.^37^ Genomic sequencing efforts revealed that plasma cfDNA in septic patients is highly digested, with fragment sizes ranging from 150 base pairs (bp) to 200 bp, with cfDNA length correlated to sepsis severity.^5,38,39^ These findings lead us to posit that unlike intact NETs and high molecular weight (HMW) DNA, it is degraded NETs and short cfDNA fragments that induce thrombosis and vascular injury, and these deleterious effects can be prevented by PF4. Here, we tested our model and investigated the mechanisms by which hPF4-mediated stabilization alters the ability of NDPs and cfDNA to trigger thrombosis and induce cellular injury.

## MATERIALS AND METHODS

### Preparation of double-stranded (ds) and single-stranded (ss) DNA

Isolated genomic DNA and NETs were digested into shorter fragments by incubating with restriction enzymes AflII, BsGrI-HF or AluI individually (10 U/μg DNA in CutSmart® buffer, all from New England BioLabs) overnight at 37°C, or DNase I (100U/mL in phosphate-buffered saline (PBS), Gibco) for 30 minutes at 37°C. ssDNA was generated by heating genomic dsDNA at 100°C for 10 min, followed by rapid freezing on dry ice to prevent reannealing. NETs and DNA fragments were size-fractionated on ethidium bromide (0.5 μg/mL) agarose gel (0.4-0.9% agarose, GeneMate) in Tris acetic acid EDTA buffer (40mM Tris, 1mM EDTA, 20mM acetic acid). Bands were visualized under UV light.^40^

### Thrombin generation assay

DNase I-digested NETs (20 μg/mL) and DNA fragments (20 μg/mL) were incubated with or without hPF4 (20 μg/mL) and KKO (20 μg/mL) in buffer containing a final concentration of 4 μM phosphatidylcholine/phosphatidylserine (75:25, Diapharma) for 10 minutes at room temperature (RT) in 96-well black MaxiSorp™ plates (475515, NUNC). Pooled normal human plasma (33% assay volume; George King Bio-Medical) was added, followed by Technothrombin® TGA substrate Z-GLY-GLY-ARG-AMC. HCl (417 μM substrate, 6.25 mM CaCl_2_ final concentration, Bachem Holding) in TGA dilution buffer (20 mM HEPES, 150 mM NaCl, 0.1% PEG-8000; pH 7.4). Fluorescence at 360nm/460nm excitation/emission (ex/em) was measured immediately and at 1-minuteintervals thereafter for 90 minutes at 37°C. Thrombin generation was calculated against a thrombin calibration curve (calibration kit, Diapharma).

### Fibrin generation assay

NETs (20 μg/mL) and DNA (20 μg/mL) were incubated with or without hPF4 (20 μg/mL) and KKO (20 μg/mL) for 10 minutes at 37°C in HBS buffer (25 mM HEPES, 150 mM NaCl; pH 7.4) in clear, medium-binding 96-well plates (9017, Corning). Pooled normal human plasma, Factor (F) XI- or FXII-depleted plasma (33% assay volume; George King Bio-Medical) was added, followed by 8.3 mM CaCl_2_ in HBS buffer. Absorbance at 405 nm was measured for 1 hour at 30-sec intervals at 37°C. Lag time (after re-calcification until first burst in fibrin generation) and slope (steepest rate of fibrin generation/min) were determined from kinetic curves. In select experiments, depleted plasma was supplemented with purified FXI (100 pM; Haematologic Technologies) or FXII (1.2 nM; Haematologic Technologies) prior to re-calcification and fibrin generation.

### Fibrinolysis assay

NETs (20 μg/mL) and dsDNA (20 μg/mL) were incubated with or without hPF4 (20 μg/mL) and KKO (20 μg/mL) prior to adding tissue-type plasminogen activator (50 μg/mL; Diapharma) in 96-well plates (Corning). Plasma was immediately added and re-calcified as described above. Absorbance at 405nm was measured for 2 hours at 1-minute intervals at 37°C. The 50% lysis time was defined as the time required to achieve a half-maximal A405 from the time maximal A405 was achieved.

### Human umbilical vein endothelial cells (HUVEC) in vitro studies

HUVECs (Lifeline Cell Technology) were grown to confluence on 0.1% gelatin (ATCC)-coated glass-bottom 8-well Ibidi® chambers with Vasculife VEGF endothelial medium complete kit (Lifeline). HUVECs were incubated with vehicle (serum-free media, SFM; Gibco), dsDNA (20 μg/mL), ssDNA (20 μg/mL), PF4 (20 μg/mL) or PF4-DNA complexes in SFM for 20 minutes at 37°C. In select experiments, HUVECs were pre-treated with toll-like receptor (TLR)-9 inhibitors E6446 (10 μM; Selleckchem) or hydroxychloroquine (HCQ) (50 μM; Selleckchem) for 30 minutes prior to incubation with vehicle, dsDNA or ssDNA fragments (20 μg/mL) in the absence or presence of PF4 (20 μg/mL) in SFM for 20 minutes at 37°C. Human α-thrombin (10 nM; Haematologic Technologies) or tumor necrosis factor alpha (TNFα, 20 ng/mL, R&D systems) were used as positive controls.

Cells were then washed with PBS and fixed in 4% paraformaldehyde (PFA) (Santa Cruz Biotechnology) for 15 minutes at RT before blocking with 3% bovine serum albumin and 2% fetal bovine serum (HyClone) in PBS for 1 hour at RT. Primary anti-human von Willebrand factor (VWF) antibody (1:1000 dilution in PBS, LOT # 75601, Agilent Dako) was incubated overnight at 4°C. Slides were washed with PBS and blocked for 1 hour at RT. Secondary Alexa Fluor-594 anti-rabbit IgG (4 μg/mL, Life Technologies) and Hoechst 33342 (Life Technologies) were incubated in PBS for 2 hours at RT in the dark. HUVECs were imaged using Zeiss 10X and 20X objective on a Zeiss LSM 710 confocal microscope. Mean fluorescence intensity (MFI) was quantified using Fiji.

HUVECs were grown to confluence in tissue culture-treated 96-well plates. Cells were washed with SFM and incubated with vehicle, digested dsDNA or ssDNA (each 20 μg/mL) and TNFα (1 ng/mL) for 16 hours in SFM in the absence or presence of PF4 (20 μg/mL). Cells were then washed with assay buffer (20 mM Tris, 100 mM NaCl, 10 mM CaCl_2_, pH 7.4). FX (160 nM, Enzyme Research Labs) and FVIIa (0.5 nM, Enzyme Research Labs) in assay buffer were added for 1 hour at 37°C. SPECTROZYME Xa (0.5 mM, Biomedica Diagnostics) was added and absorbance at 405 nm was immediately recorded for 1 hour at 1-minute intervals to determine maximum rate of reaction, V_max_ (mOD/min).

### LPS endotoxemia model

Wild-type (WT) C57BL/6 mice received an intraperitoneal injection of LPS (35 mg/kg). A subset of animals was given 40 mg/kg of hPF4 by tail-vein injection immediately following LPS injection. Six hours following LPS injection, animals were euthanized, blood was collected from the inferior vena cava into 3.2% sodium citrate (1:10 v/v), and platelet-poor plasma was isolated by centrifugation at 2000g for 20 minutes at RT.

Thrombin anti-thrombin complexes in septic plasma were measured using commercially available ELISA kits (AssayPro; EMT1020-1) according to manufacturers’ instructions. Briefly, citrated plasma was diluted 125-fold and incubated with polyclonal antibody against mouse thrombin for 2 hours at RT. 96-well plates were washed and incubated with biotinylated mouse TAT complex antibody for 1 hour at RT. The microplate was then incubated with streptavidin-peroxidase conjugate for 30 minutes at RT, and chromogenic substrate was then added for 5 minutes at RT. Reactions were stopped and absorbance was measured at 450nm immediately.

Plasma from LPS-treated mice were diluted 1:10 with Hanks’ Balanced Salt Solution (no Ca^+2^/Mg^+2^; Gibco) in 96-well black MaxiSorp™ plate. Equal volume of SYTOX Green (5 μM final in HBSS without Ca^+2^/Mg^+2^, Invitrogen) was added to each sample and incubated in the dark for 10 min. Fluorescence at 485/527 ex/em was measured at RT, and plasma cfDNA concentrations were determined based on fluorescence of calf thymus DNA curve of 0 to 10 μg/mL.

### Statistical analyses

Data are presented as mean ± standard error of the mean (SEM). The Shapiro-Wilk normality test was used to determine whether group data were distributed normally. Levene’s test was used to determine equality of variances. One-way ANOVA with Tukey’s post hoc test was used to compare between treatment groups. Mann-Whitney test or Kruskal-Wallis with Dunn post hoc test was used to compare between groups when the data did not qualify for parametric statistics. *P* ≤ 0.05 was considered significant. All statistical analyses were conducted using GraphPad Prism 9.

### Study ethics approval

Human blood for studies was collected after informed consent from healthy, aspirin-free volunteers using a 19-gauge butterfly needle in 3.8% sodium citrate (10:1 v/v) under a protocol approved by the Children’s Hospital of Philadelphia (CHOP) Institutional Review Board and were consistent with the Helsinki Principles. Animal procedures were approved by the Institutional Animal Care and Use Committee (IACUC) in accordance with NIH guidelines and the Animal Welfare Act.

### Supplemental methods and materials

See the Supplement Materials and Methods for further details on reagents used, isolation of human neutrophils, preparation of NETs, isolation of genomic DNA from human white blood cells, as well as the details on the intravenous administration of DNA into mice.

### Data sharing statement

For original data, please contact.

## RESULTS

### PF4 and KKO inhibit fibrin generation initiated by NET and DNA fragments

We treated intact NETs and HMW DNA of >50 kb (Figure S1A) with DNase I to generate NDPs and low molecular weight (LMW) DNA measuring 0.1-0.5 kb (Figure S1B). To determine whether DNA fragment length influences NET-mediated fibrin generation, HMW DNA (>50 kb) was digested with six-cutter restriction enzymes, AflII and BsGrI-HF, and the 4-cutter, AluI, to generate DNA fragment lengths of ~4 kb and ~250 bp, respectively (Figure S1C). We then assessed the thrombogenicity of NET and DNA fragments in plasma using a fibrin-generation assay. HMW NETs and genomic DNA did not induce fibrin generation in plasma (Figure 1A); however, digested NETs shortened lag time from ~900 to ~720 seconds (Figure 1A), while DNA fragments of various lengths cleaved by restriction enzymes all significantly shortened lag time to ~650 seconds (Figure 1A) and increased the rate (slope) of fibrin generation (Figure 1B). hPF4 at 20 μg/mL, a concentration achieved at sites of thrombus formation ^41^, reversed DNA-induced, fibrin-generation lag time to baseline, regardless of chain length (Figure 1C).

**Figure 1.**
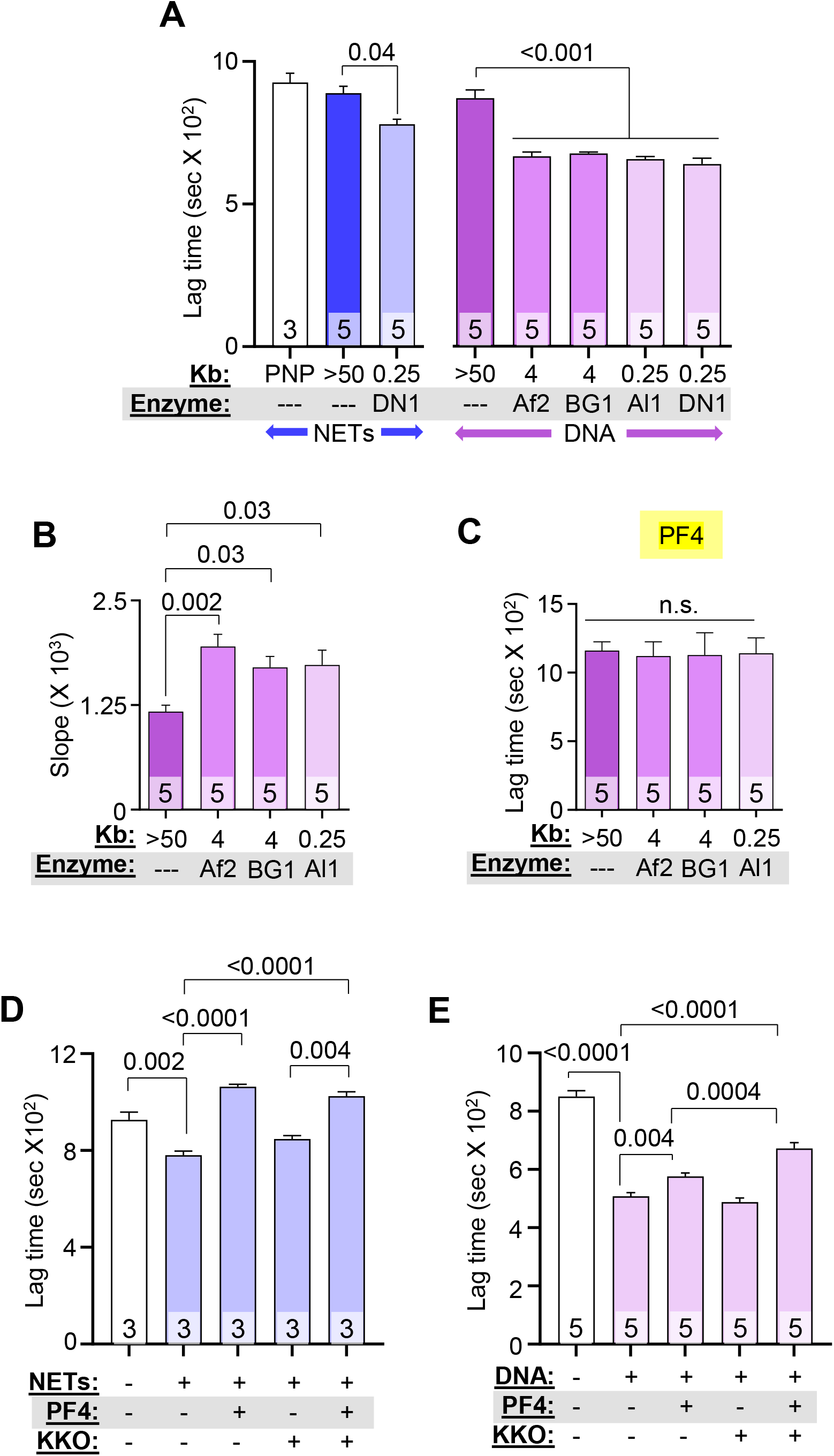
PF4 and KKO inhibits fibrin generation initiated by NET and DNA fragments. Fibrin generation in pooled normal plasma (PNP) with or without added HMW, digested NETs (blue), or DNA (purple). DN1 = DNase I, Af2 = AflII, BsGrI-HF = BG1 and Al1 = AluI. Data are mean ± SEM) of at least 3 independent experiments as indicated in each bar. (**A)** Lag time determined from kinetic curves of fibrin generation with NETs (blue) and DNA (purple). (**B**) Slope (rate) of fibrin generation was determined from the same kinetic curves as in (A). (**C**) Lag time of DNA-induced fibrin generation was determined from kinetic curves similarly to (A), in the presence of PF4 (20 μg/mL). (**D-E**) Lag times are shown for fibrin generation studies of DNase I-digested NETs (**D**) and DNA (**E**) in the absence or presence of 1 μg/mL low PF4 and/or 10 μg/mL KKO. Data are mean ± SEM of at least 3 independent experiments as indicated in each bar.

Although we previously observed that binding of KKO to hPF4:NET complexes increased the resistance of NETs to nuclease digestion,^31^ inclusion of KKO did not alter the anticoagulant effect of PF4 on digested DNA at the studied concentration (Figure S2A). At a lower concentration (1 μg/mL), hPF4 still fully reversed the prothrombotic effects of NETs (Figure 1D), but was only partially effective in the case of isolated DNA (Figure 1E). This difference between NETs and DNA could be due to the presence of positively-charged histones bound to the polyphosphate backbone of NETs. The addition of KKO (10 μg/mL) to DNA significantly prolonged fibrin lag time in the presence of low-dose hPF4 (Figures 1E and S2B). These results indicate that hPF4 complexed to either NETs or DNA reduced their fibrin generation potential in human plasma.

### hPF4 and KKO inhibit thrombin generation, but not fibrinolysis, by NET and DNA fragments

We next measured thrombin generation to address the mechanism by which hPF4 and KKO inhibited the procoagulant effects of NETs and DNA. hPF4 significantly reduced NET-induced peak thrombin generation and delayed thrombin-initiation time (Figures 2A and 2B). hPF4 also significantly reduced DNA-induced thrombin generation and delayed thrombin lag time (Figures 2A and 2B). The addition of KKO did not alter hPF4-mediated effects on NET- or DNA-induced thrombin generation (Figures 2A and 2B, and Figure S3A).

**Figure 2.**
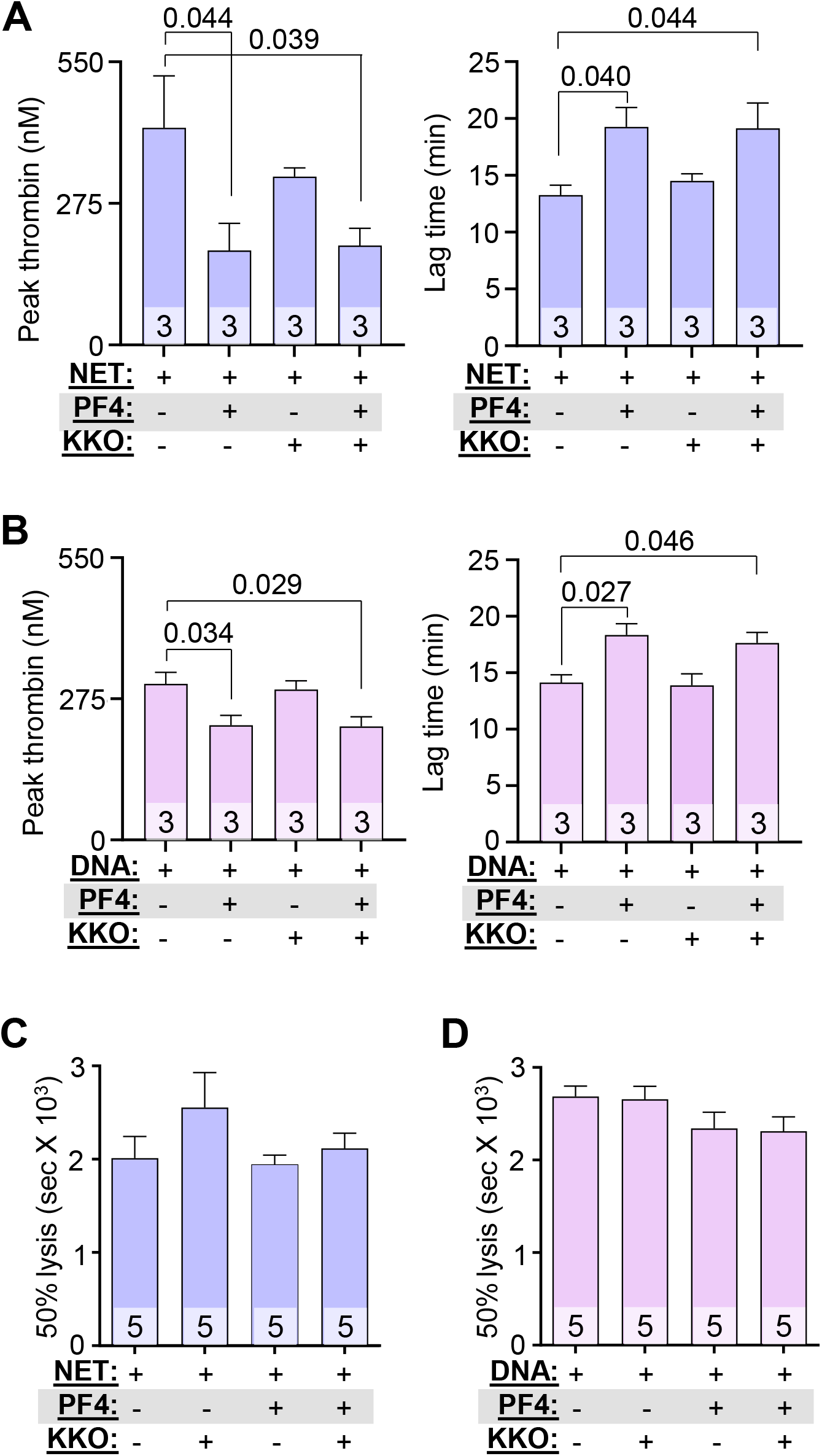
PF4 and KKO inhibit thrombin generation but not fibrinolysis. (**A** and **B**) Thrombin generation studies of DNase-digested NETs (**A**) and DNA (**B**) in the absence or presence of PF4 (20 μg/mL) and/or KKO (10 μg/mL). (**C** and **D**) Time to 50% clot lysis of NETs- (**C**) and DNA- (**D**) induced coagulation, as determined based on kinetic curves. Data are mean ± SEM of at least 3 independent experiments as indicated in each bar.

There have been conflicting reports concerning the effect of NETs and cfDNA on thrombus stability. Some studies show NETs potentiate fibrinolysis by stimulating fibrin-independent plasminogen activation, while others observe that NET components such as neutrophil elastase and cfDNA promote resistance to thrombolysis by degrading plasminogen and forming lysis-resistant complexes with fibrin, respectively.^42,43^ Therefore, we assessed whether hPF4:NET or hPF4:DNA complexes affect the rate of internal fibrinolysis in NETs- and DNA-induced plasma clots by mixing a plasminogen activator with NETs or DNA with or without added hPF4 prior to re-calcification. While hPF4 (1 and 20 μg/mL) and KKO (10 μg/mL) delayed NETs- and DNA-induced fibrin generation significantly as described above, we observed no difference in time to clot lysis (Figure 2C and 2D). This indicates that hPF4, which prevents DNases from digesting NETs,^31^ does not interfere with fibrinolysis.

### hPF4 modulates fibrin formation through FXI and XII

We next sought to define the mechanism by which hPF4 modulates the prothrombotic effects of cfDNA. Polyphosphates like DNA trigger thrombin generation in plasma by activating the intrinsic pathway of coagulation, specifically FXI and FXII.^44–47^ In fibrin-generation studies using FXI- and FXII-depleted plasma, hPF4, at concentrations as high as 20 μg/ml, had no effect on DNA-induced fibrin generation (Figure 3A and Figure S3B). Upon supplementation of FXI-depleted plasma with FXI, hPF4 significantly delayed DNA-induced fibrin generation lag time (Figure 3B and Figure S4A). hPF4 also delayed fibrin-generation in FXII-depleted plasma supplemented with FXII (Figure 3B and Figure S4B). These results suggest that hPF4 blocks the procoagulant activity of DNA in plasma by interfering with DNA-mediated intrinsic coagulation pathway activation.

**Figure 3.**
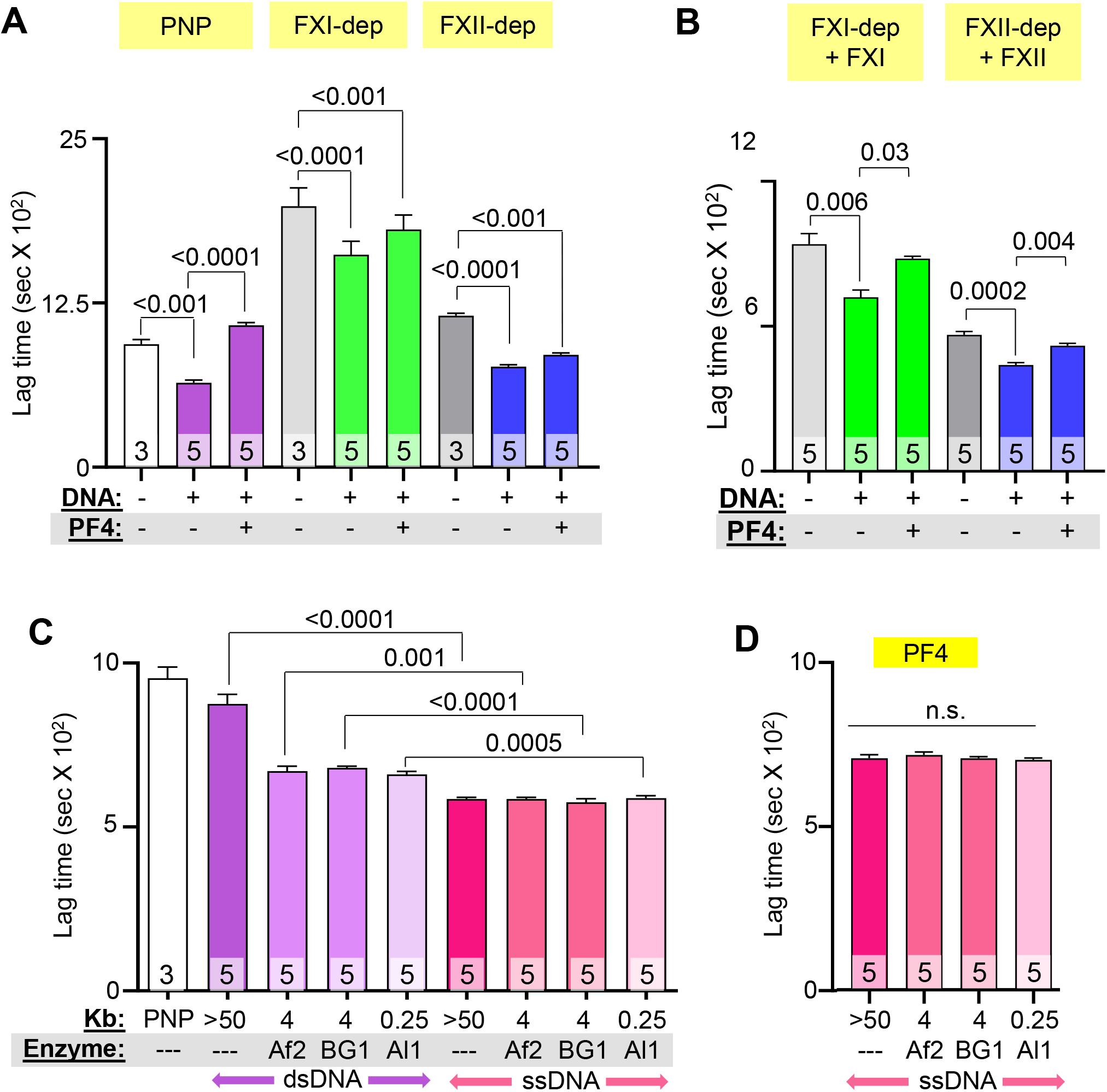
Effect of PF4 on fibrin generation in FXI- and FXII-depleted plasma and on thrombogenicity of ssDNA. (**A**) Lag time, determined from kinetic curves, of fibrin generation in PNP or FXI- or FXII-depleted (dep) plasma, induced by digested DNA in the absence or presence of PF4. (**B**) Lag time of fibrin generation induced by digested DNA in depleted plasma supplemented with missing coagulation factors. (**C**) Lag time of ssDNA-induced (pink) fibrin generation compared to dsDNA (purple). (**D**) Lag time of ssDNA-induced fibrin generation with added PF4 (20 μg/mL). Data are mean ± SEM of at least 3 independent experiments as indicated in each bar.

### hPF4 attenuates the thrombogenicity of ssDNA

We hypothesized that shorter NETs and DNA fragments are more thrombogenic than HMW NETs and genomic DNA because they exhibit greater levels of ssDNA at their termini and form hairpin structures that are potent activators of coagulation.^48^ To investigate this hypothesis, we denatured genomic DNA fragments of different size ranges at 100°C for 10 minutes and then rapidly froze the samples to generate ssDNA of various lengths. ssDNA fragments of all lengths dramatically accelerated fibrin generation, an interval significantly shorter than lag times seen with HMW and LMW dsDNA (Figure 3C). At the same concentration, hPF4 delayed fibrin lag time by ssDNA, but not to the same extent as seen with dsDNA (Figure 3D versus Figure 3C), perhaps because ssDNA is much more thrombogenic than dsDNA.

### hPF4 protects endothelium from activation of prothrombotic pathways by DNA

Our group previously found that hPF4 decreases NET-mediated endothelial toxicity.^31^ Building on this observation, we assessed whether PF4 modulates the ability of digested DNA to promote a procoagulant phenotype in endothelial cells. HUVECs were incubated with DNA fragments of different size ranges with and without hPF4, prior to staining for VWF release as an indicator of HUVEC activation. HMW DNA did not induce VWF secretion by HUVECs, while short dsDNA fragments (0.25-4kb) significantly increased surface VWF levels similar to that seen upon thrombin activation^49^ (Figure 4A and 4B, and Figure S5A). PF4 prevented dsDNA-induced VWF secretion regardless of DNA length (Figure 4A and Figure S5A). Prior studies showed that internalized unmethylated CpG motifs in bacterial DNA activate endothelial cells via TLR9.^50^ We now treated cultured HUVECs with the TLR9 inhibitors E6446 and hydroxychloroquine (HCQ), and found that both TLR9 inhibitors abolished dsDNA-induced VWF secretion (Figure 4B and Figure S5C).

**Figure 4.**
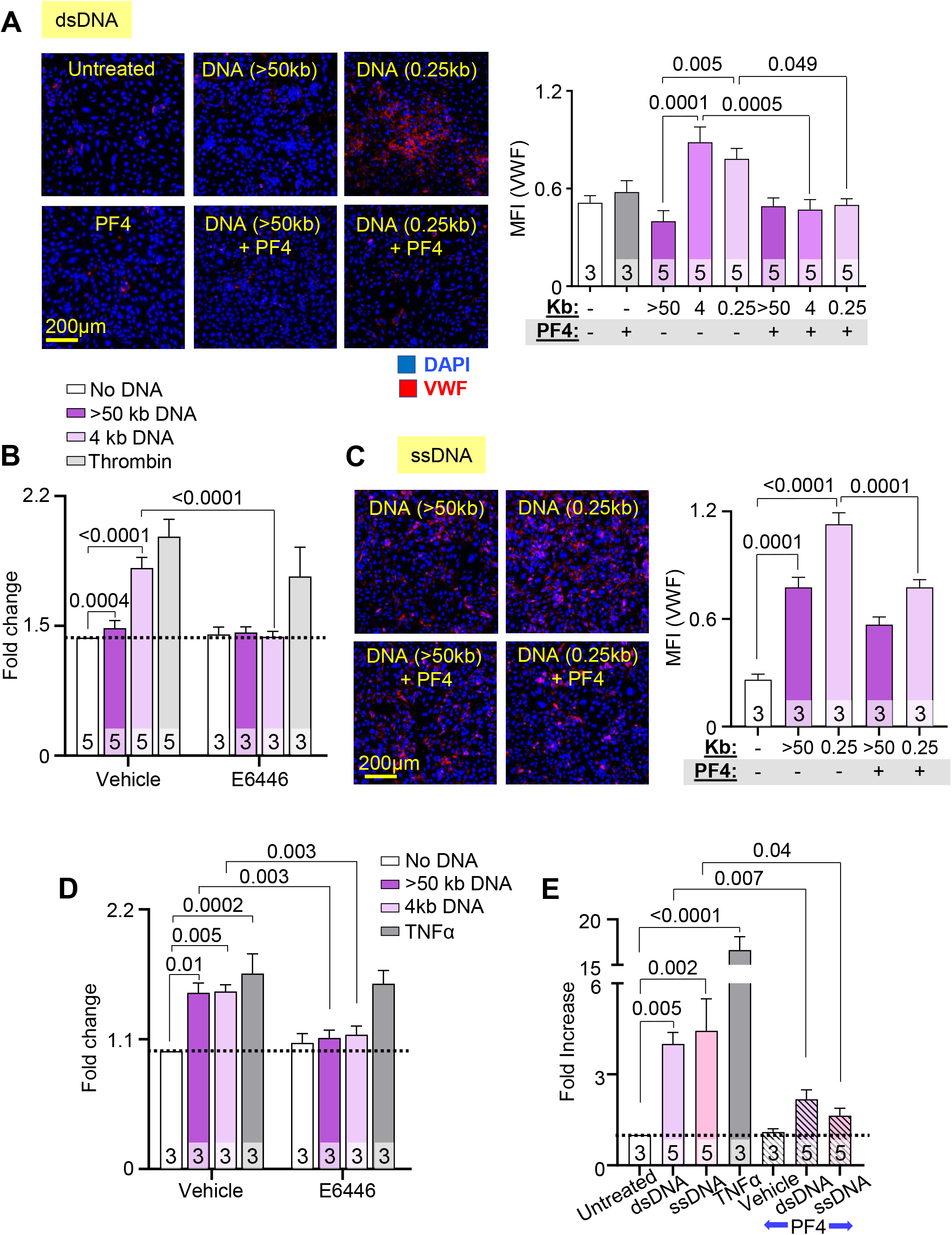
PF4 protects against dsDNA and ssDNA-induced procoagulant responses by endothelium. Mean fluorescence intensity (MFI) of released VWF from HUVECs exposed to fragments of dsDNA (**A** and **B**) or ssDNA (**C** and **D**), with or without PF4 or TLR9 inhibitor (E6446). Data were normalized to MFI of untreated cells without exposure to dsDNA or inhibitor (as indicated by dotted lines). Exposure to α-thrombin (**B**) or TNFα (**D**) serves as positive controls. (**E**) TF expression by HUVEC (shown as fold increase from untreated cells without PF4 exposure) induced by dsDNA or ssDNA in the absence or presence of PF4. TNFα exposure was used as a positive control. Data are mean ± SEM of at least 3 independent experiments as indicated in each bar.

Unlike HMW dsDNA, HMW ssDNA stimulated robust VWF release comparable to levels of release seen with LMW dsDNA. ssDNA fragments induced VWF release at nearly twice the level seen with similar-sized dsDNA and comparable to that seen using exposure to TNFα^51^ (Figures 4C, 4D and Figure S5B). hPF4 significantly reduced endothelial release of VWF by ssDNA, although not as effectively as after exposure to dsDNA (Figure 4C). Inhibition of TLR9 with E6446, and to a lesser extent HCQ, reduced ssDNA-induced VWF secretion (Figure 4D and Figure S5D).

NETs induce endothelial TF expression,^15^ but it remains unclear which components induce this effect. To determine whether NET cfDNA up-regulates endothelial TF expression, HUVECs were incubated with LMW dsDNA and ssDNA fragments overnight prior to exposure to FX and activated FVII (FVIIa). Rate of cleavage of a chromogenic substrate by activated FX (FXa) was measured as an indicator of DNA-induced endothelial TF expression. Both LMW dsDNA and ssDNA fragments enhanced TF expression ~4.5 fold compared to vehicle alone, while the inclusion of hPF4 during overnight incubation significantly reduced endothelial TF expression induced by both dsDNA- and ssDNA (Figure 4E).

### cfDNA increases plasma TAT levels in Cxcl4^-/-^ mice

To assess thrombogenicity of cfDNA *in vivo* and to define the role of PF4 in modulating DNA-induced thrombosis, WT or *cxcl4^-/-^* littermate mice were given an intravenous bolus of HMW DNA, and plasma levels of TAT complexes were analyzed. Although there was no difference in TAT levels between WT and *cxcl4^-/-^* mice 30 minutes post-DNA infusion, *cxcl4^-/-^* mice had significantly elevated TAT levels compared to WT mice 4 hours following DNA injection (Figure 5A).

**Figure 5.**
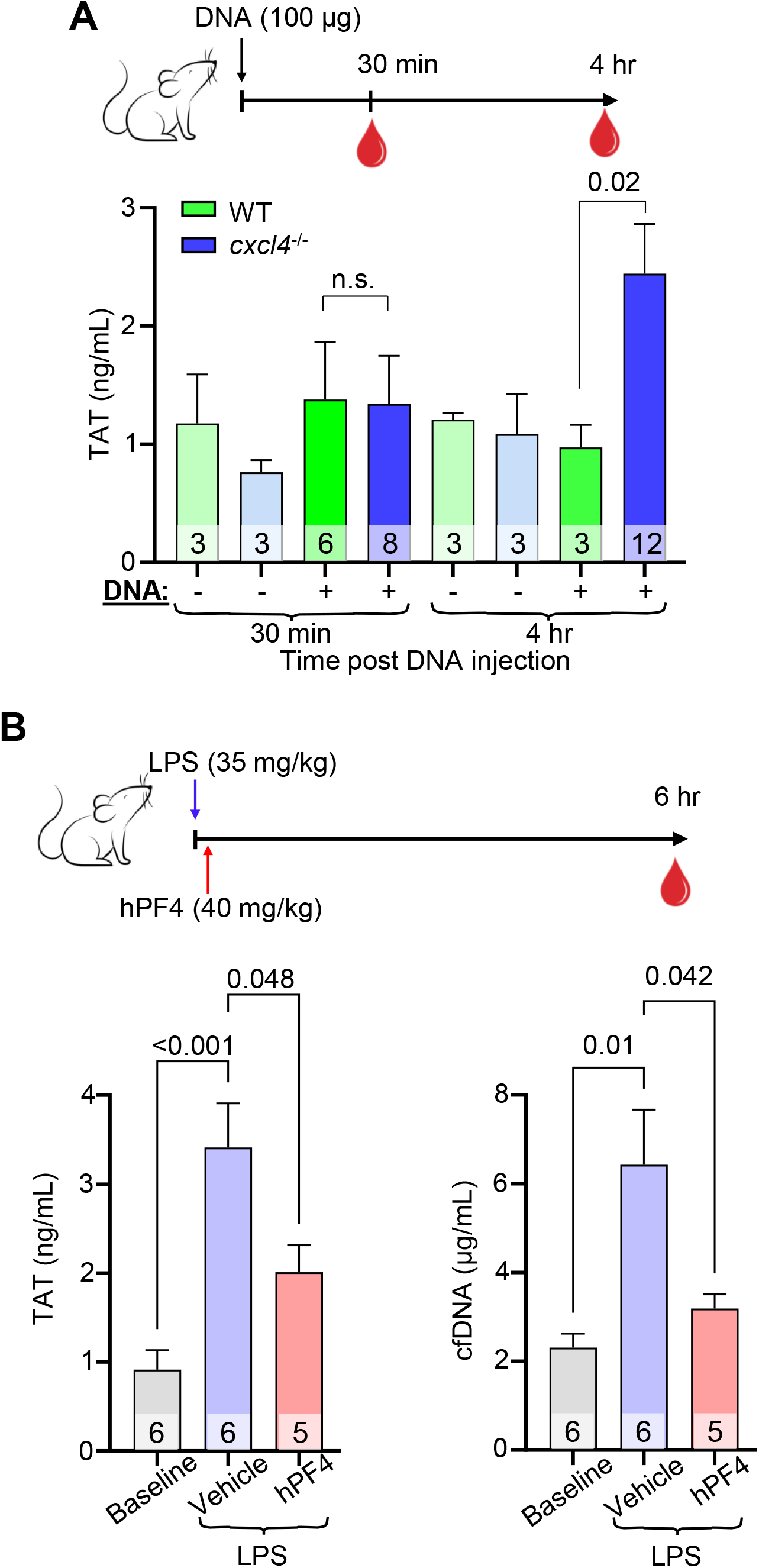
PF4 attenuates cfDNA-induced coagulability in vivo. (**A**) Top: schematic of study in which WT mice or *cxcl4^-/-^* littermates were given normal saline vehicle or normal saline containing digested DNA prior to blood collection at 30 minutes or 4 hours. Bar graph: TAT levels using a commercial ELISA kit in platelet-poor plasma. Data are mean ± SEM of at least 3 independent experiments as indicated in each bar. (**B**) Top: schematic of study in which WT mice received normal saline vehicle or LPS, and a subset of animals was given hPF4 by tail-vein injection immediately following LPS injection. TAT levels were measured in blood was drawn at baseline and 6-hours post LPS ± PF4 infusion. Bar graphs: TAT levels (left) and cfDNA levels (right) at 6-hours post LPS challenge. Data are mean ± SEM of at least 5 independent experiments as indicated in each bar.

### hPF4 inhibits generation of TAT by cfDNA in endotoxemic mice

To determine if PF4 modulates the thrombogenicity of cfDNA released during sepsis, WT mice were given an intraperitoneal injection of LPS to induce endotoxemia, an intervention shown by our group and others to induce a significant rise in plasma cfDNA.^22,31^ A subset of these mice was given vehicle or hPF4 by tail-vein injection immediately following LPS injection. Six hours post-LPS injection, saline-treated mice exhibited a significant elevation in TAT levels as compared to baseline (Figure 5B). In contrast, mice administered a single bolus of hPF4 had significantly reduced TAT (Figure 5B, left) and cfDNA (Figure 5B, right) levels.

## DISCUSSION

It may seem counterintuitive to propose NET stabilization as a therapeutic strategy, as elevated NET markers have been associated with increased disease severity in a diverse range of thromboinflammatory disorders including sepsis, COVID-19 and sickle cell disease.^16,17,52–72^ However, studies from multiple groups show that the formation of NETs is an evolutionarily conserved function, initially developed in protozoa and retained as part of the innate immune response of plants and animals to capture invading pathogens and immobilize them near concentrated antimicrobial compounds.^73^ Intact NETs are not pathogenic when released in a regulated manner, and indeed may limit collateral host tissue damage by preventing the systemic release of toxic NDPs, including cfDNA (Figure 6A).

**Figure 6.**
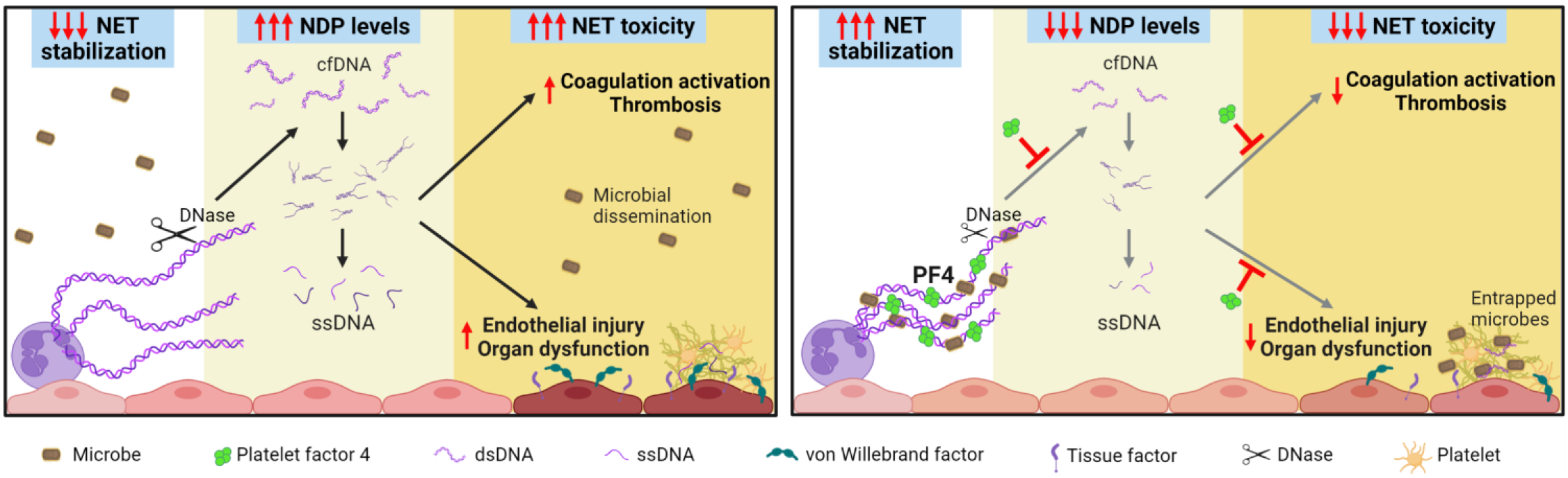
Proposed protective mechanisms of NET-stabilization. **Left**: NETs are subjected to digestion by DNase I, reducing microbial capture and liberating toxic NDPs, including cfDNA. These cfDNA fragments expose ssDNA at termini that trigger coagulation activation and induce endothelial injury, leading to thrombosis and end-organ dysfunction in sepsis. **Right**: NET stabilization by PF4 enhances DNase I resistance, enhancing NET microbial capture, reducing circulating cfDNA levels, and attenuating cfDNA-induced thrombogenicity and toxicity to endothelial cells.

To date, NET-targeted strategies proposed for the treatment of sepsis have fallen into two categories: those that prevent NETosis^21,74–85^ and those that accelerate NET degradation.^86–97^ Neither strategy enhances the beneficial effects of NETs; the latter strategy may also suffer from liberating captured pathogens and increasing circulating levels of NDPs. There remains an unmet clinical need for a non-injurious therapeutic approach in sepsis. Our group previously showed that hPF4 binds to NETs, causing them to become physically compact and resistant to nucleases, limiting the release of NDPs.^31^ Moreover, the positively-charged PF4 tetramer, found in high molar concentrations at sites of platelet degranulation, cross-binds negatively-charged microbes to the negatively-charged DNA scaffold of NETs. Without PF4, microbial entrapment may be markedly impaired.^31^ That study showed that PF4 has positive effects on NETs, preventing excessive NET degradation and enhancing the protective effect of NETs while minimizing their capacity to induce harm.

We posited that smaller fragments of dsDNA have a higher thrombotic potential because they have a greater number of termini that can “breathe”, exposing ssDNA^98^. To explore this idea, we generated ssDNA by melting HMW genomic DNA and observed that irrespective of fragment length, ssDNA was more prothrombotic and injurious to the endothelium than dsDNA. It is likely that shorter dsDNA fragments, cleaved from digested NETs, exhibit higher DNA end-breathing, exposing more ssDNA and hairpin loops and avidly bind kininogen, serving as potent activators of the intrinsic coagulation pathway.^48^ Moreover, ssDNA internalized by endothelial cells binds to TLR9 to induce a prothrombotic phenotype.^99^ Others have shown that DNA binding to proteins present in high concentrations in NETs, including HMGB1 and LL37, enhance DNA uptake and TLR9 activation.^100^

The PF4 tetramer may be uniquely able to combat the prothrombotic effects of cfDNA because it has a positively-charged circumferential zone that can bind to two polyanions, one on each side.^101^ This bivalent property enables a single tetramer of PF4 to simultaneously bind the polyphosphate-ribose backbone of two separate DNA chains, leading to NET compaction and cfDNA aggregation. Our in vitro studies show that PF4, at concentrations that are well within the range seen in the setting of thromboinflammation,^41^ can reduce the prothrombotic and endothelial-activating properties of LMW dsDNA and ssDNA. Our in vivo experiments in mice infused with genomic DNA or treated with LPS, similarly demonstrate that the expression of PF4 protects animals from plasma cfDNA-enhanced thrombosis, consistent with our model (Figure 6B) and our prior studies that showed treatment with hPF4 improved outcomes in murine sepsis.^31^

Of note, further stabilization of hPF4:NET complexes with KKO, does not interfere with the protective effects of PF4 on procoagulant activity and endothelial cell activation or impair thrombolysis. Indeed, hPF4 and DGKKO likely work in concert to reduce the pathogenic effects of cfDNA.^31^ Because platelet counts and hPF4 content per platelet vary in the general population,^102,103^ and critically ill individuals may have increased platelet activation with high plasma PF4 levels or thrombocytopenia,^104,105^ septic patients may benefit from either hPF4 and/or DG-KKO infusion to optimize NET stabilization.

In summary, NETs have an important positive role in host defense, impeding the spread of pathogens and limiting the systemic release toxic antimicrobial compounds; however, when rapidly degraded, NETs release cfDNA that can initiate the contact pathway of coagulation and activate the endothelium in part via TLR9. We previously showed that PF4 enhances NET microbial entrapment, and now show that it limits the release of LMW dsDNA and ssDNA, decreases the procoagulant effects of NETs and prevents endothelial injury. DGKKO, which further decreases NET susceptibility to nucleases, does not interfere with PF4’s ability to prevent pathologic thrombosis and maintain endothelial integrity. These studies support further investigation of NET stabilization strategies to enhance the protective effect of NETs, while limiting the toxicities of plasma cfDNA.

## AUTHOR CONTRIBUTIONS

A.T.P. Ngo contributed to experimental design, performed experiments, analyzed and evaluated data and wrote the manuscript. A. Sarkar, N. Levine, V. Bochenek and I. Yarovoi performed experiments and edited the manuscript. G. Zhao, L. Rauova, M. A. Kowalska, K. Eckart, N. Mangalmurti, A. Rux contributed to mouse breeding, experimental design and data interpretation. D. Cines contributed to discussions of the proposed research and to editing the manuscript. M. Poncz and K. Gollomp conceived, designed, supervised the project and contributed to writing of the manuscript.

