## Supplemental figures for "Neutrophil extracellular trap stabilization by platelet factor 4 reduces thrombogenicity and endothelial cell injury"

### Slide 1
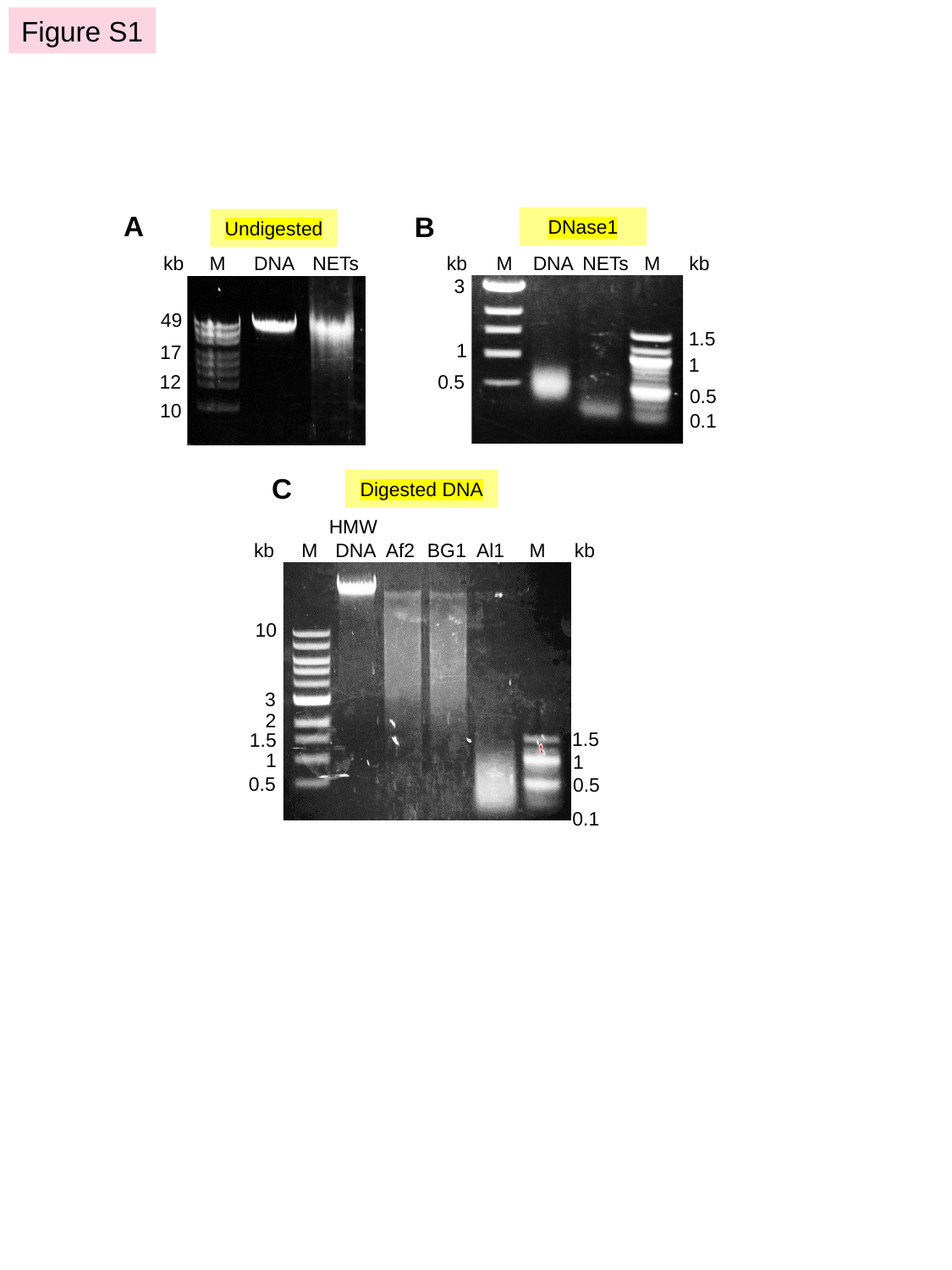

Figure S1
A
B
DNase1
Undigested
kb
M
DNA
NETs
M
kb
3
1.5
1
1
0.5
0.5
0.1
M
DNA
NETs
kb
49
17
12
10
C
Digested DNA
HMW
DNA
kb
M
Af2
BG1
Al1
M
kb
10
3
2
1.5
1.5
1
1
0.5
0.5
0.1

### Slide 2
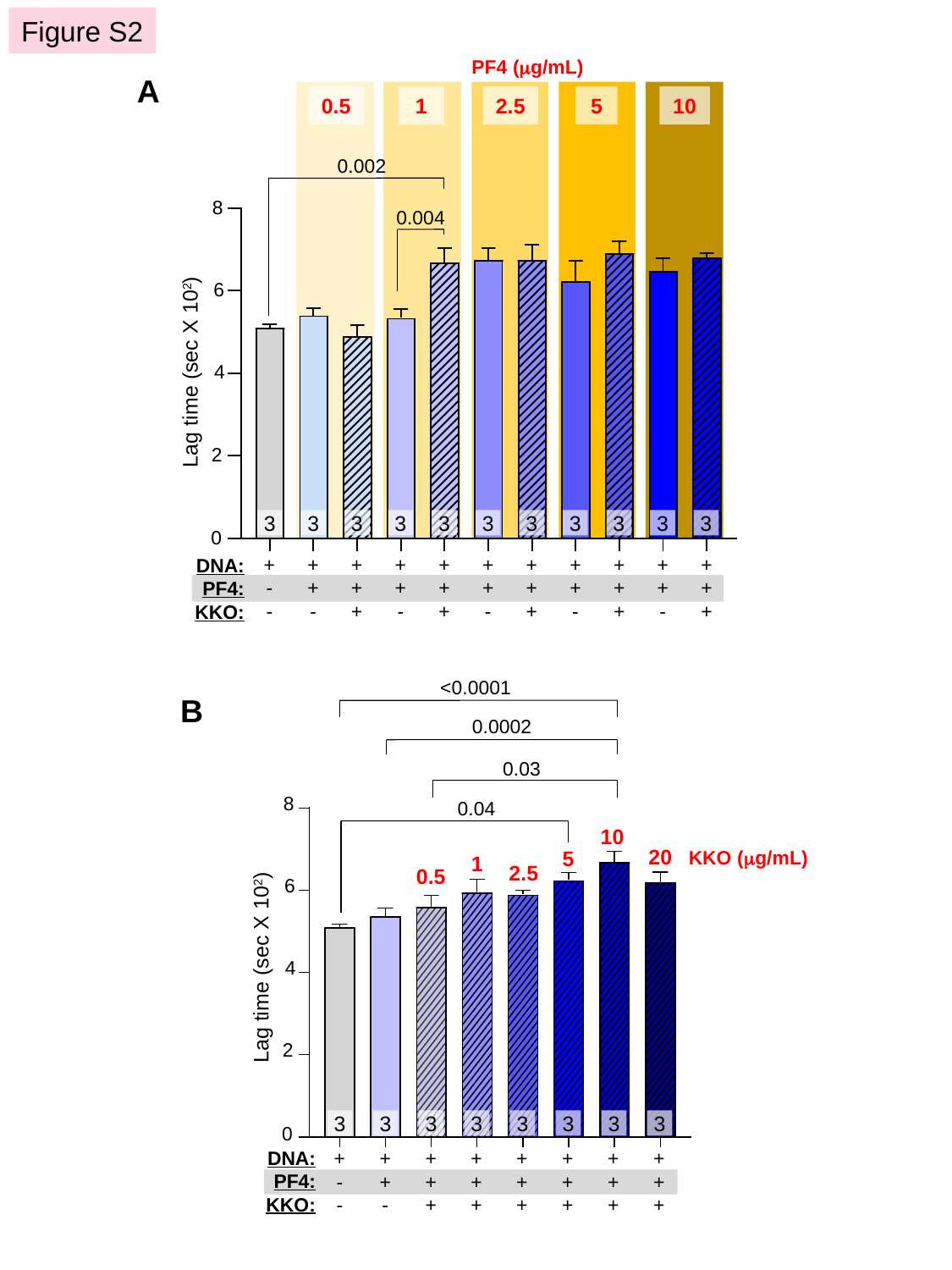

Figure S2
PF4 (μg/mL)
A
0.5
1
2.5
5
10
0.002
8
0.004
6
Lag time (sec X 10^2^)
4
2
3
3
3
3
3
3
3
3
3
3
3
0
+
-
-
+
+
-
+
+
+
+
+
-
+
+
+
+
+
-
+
+
+
+
+
-
+
+
+
+
+
-
+
+
+
DNA:
PF4:
KKO:
<0.0001
B
0.0002
0.03
8
0.04
10
20
KKO (μg/mL)
5
1
2.5
0.5
6
Lag time (sec X 10^2^)
4
2
3
3
3
3
3
3
3
3
0
DNA:
PF4:
KKO:
+
-
-
+
+
-
+
+
+
+
+
+
+
+
+
+
+
+
+
+
+
+
+
+

### Slide 3
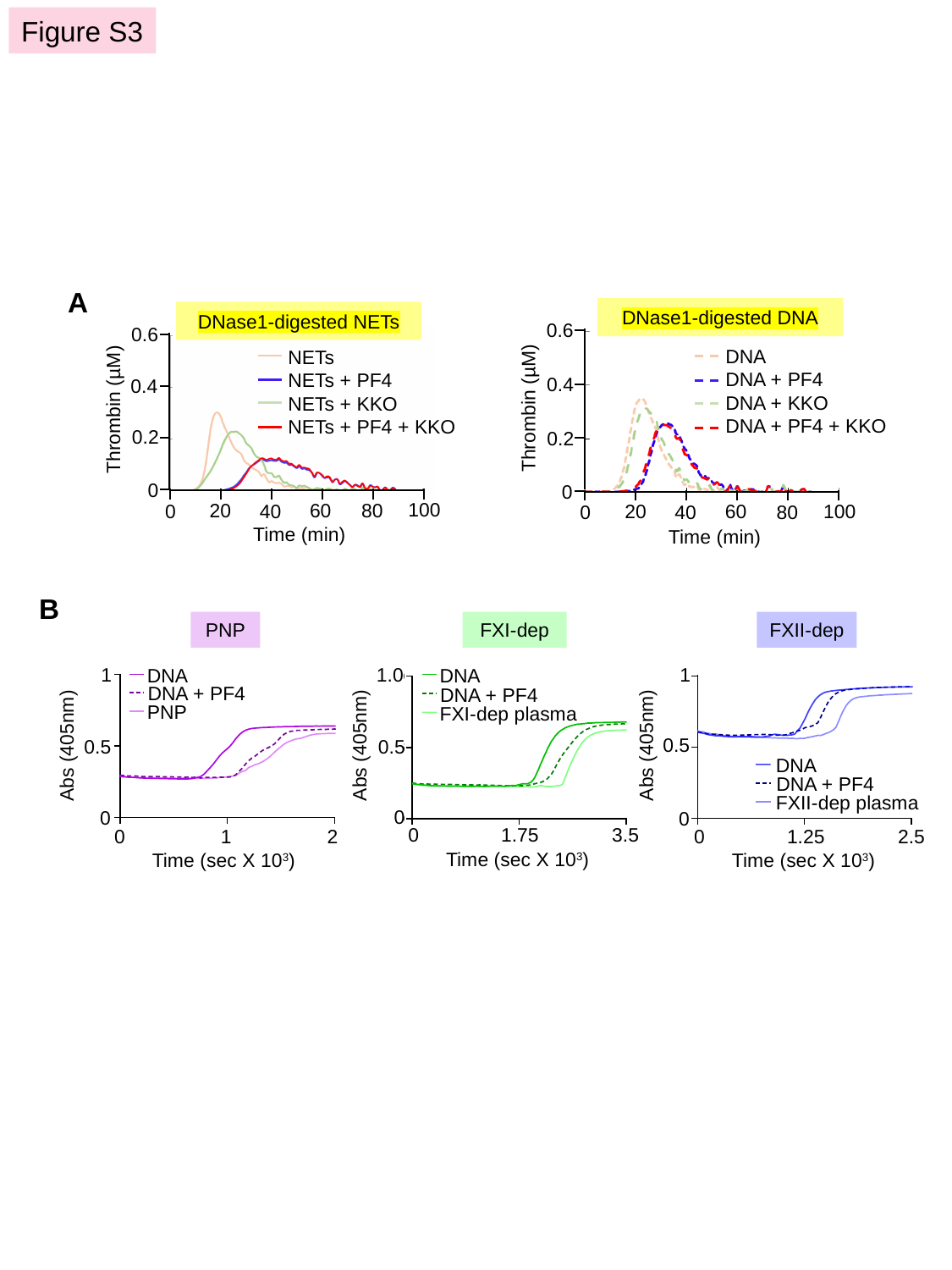

Figure S3
A
DNase1-digested DNA
DNase1-digested NETs
0.6
0.4
Thrombin (µM)
0.2
0.6
0.4
Thrombin (µM)
0.2
DNA
DNA + PF4
DNA + KKO
DNA + PF4 + KKO
NETs
NETs + PF4
NETs + KKO
NETs + PF4 + KKO
0
0
100
20
60
80
0
40
100
20
60
0
80
40
Time (min)
Time (min)
B
PNP
FXI-dep
FXII-dep
1.0
1
1
DNA
DNA
DNA + PF4
DNA + PF4
PNP
FXI-dep plasma
0.5
Abs (405nm)
Abs (405nm)
Abs (405nm)
0.5
0.5
DNA
DNA + PF4
FXII-dep plasma
0
0
0
0
1.75
3.5
Time (sec X 10^3^)
0
1
2
Time (sec X 10^3^)
0
1.25
2.5
Time (sec X 10^3^)

### Slide 4
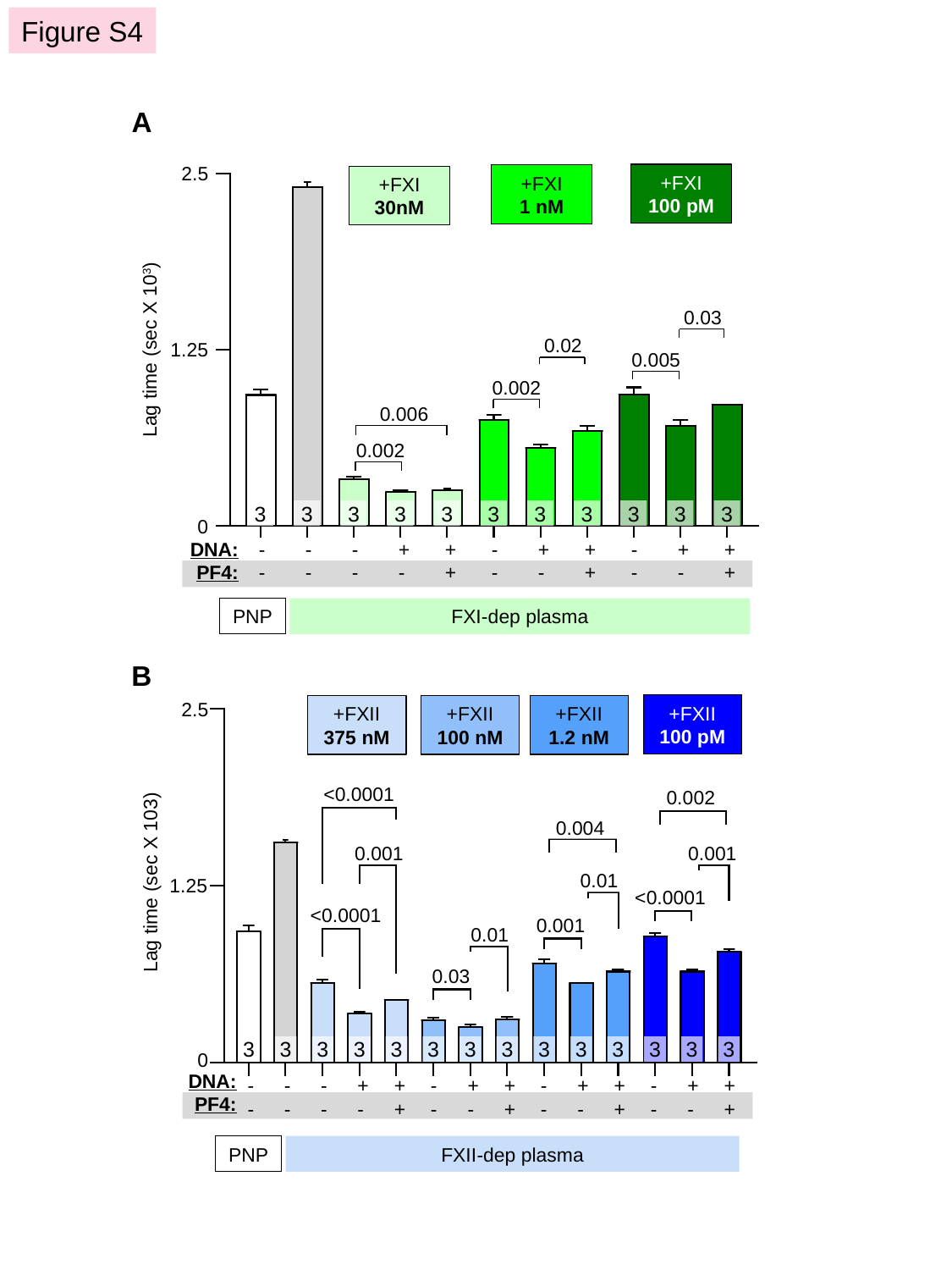

Figure S4
A
2.5
+FXI
100 pM
+FXI
1 nM
+FXI
30nM
0.03
0.02
Lag time (sec X 10^3^)
1.25
0.005
0.002
0.006
0.002
3
3
3
3
3
3
3
3
3
3
3
0
-
-
-
-
-
-
+-
++
-
-
+-
++
-
-
+-
++
DNA:
PF4:
PNP
FXI-dep plasma
B
2.5
+FXII
100 pM
+FXII
375 nM
+FXII
100 nM
+FXII
1.2 nM
<0.0001
0.002
0.004
0.001
0.001
0.01
Lag time (sec X 103)
1.25
<0.0001
<0.0001
0.001
0.01
0.03
3
3
3
3
3
3
3
3
3
3
3
3
3
3
0
DNA:
PF4:
-
-
-
-
-
-
+-
++
-
-
+-
++
-
-
+-
++
-
-
+-
++
PNP
FXII-dep plasma

### Slide 5
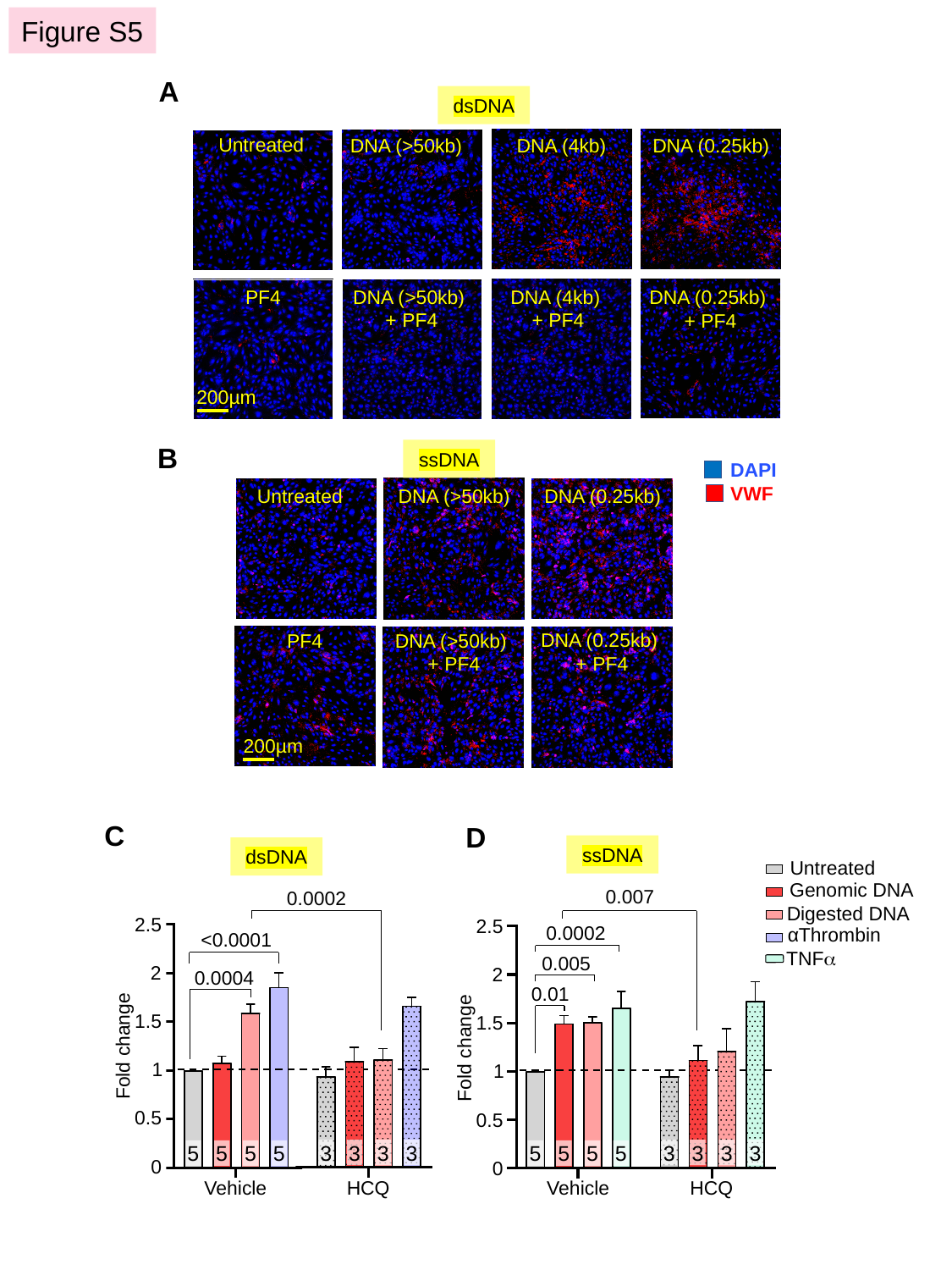

Figure S5
A
dsDNA
Untreated
DNA (>50kb)
DNA (4kb)
DNA (0.25kb)
PF4
DNA (>50kb)
+ PF4
DNA (4kb)
+ PF4
DNA (0.25kb)
+ PF4
200µm
B
ssDNA
DNA (0.25kb)
Untreated
DNA (>50kb)
DNA (0.25kb)
+ PF4
PF4
DNA (>50kb)
+ PF4
200µm
DAPI
VWF
C
D
ssDNA
dsDNA
Untreated
Genomic DNA
0.007
0.0002
Digested DNA
2.5
2.5
αThrombin
0.0002
<0.0001
TNFα
0.005
2
2
0.0004
0.01
1.5
1.5
Fold change
Fold change
1
1
0.5
0.5
3
3
3
3
3
3
3
3
5
5
5
5
5
5
5
5
0
0
Vehicle
HCQ
Vehicle
HCQ
