## Supplement for "Neutrophil extracellular trap stabilization by platelet factor 4 reduces thrombogenicity and endothelial cell injury"

#### SUPPLEMENTAL MATERIALS AND METHODS

*Reagents*

All reagents were purchased from Sigma-Aldrich except as noted. Recombinant hPF4 was expressed in S2 cells and purified using affinity chromatography and protein liquid chromatography as previously described[1]. The end-product was found to be endotoxin-free using ToxinSensor™ chromogenic LAL endotoxin assay kit (Genscript), and was tested for size distribution by SDS-polyacrylamide gel electrophoresis. KKO is a mouse $IgG_{2b\kappa}$ anti-hPF4-heparin monoclonal antibody, purified from hybridoma supernatants[2]. DNA 1kb and 100bp ladders were from New England BioLabs. GeneRuler high range DNA ladder was from ThermoFisher.

*Human neutrophil isolation and NET preparation*

Human whole blood collected into 3.8% sodium citrate (1:10 v/v) was layered over an equal volume of Lympholyte®-Poly density gradient (CedarLane Labs) and centrifuged at 500g for 35 minutes at RT. The neutrophil layer was collected and washed with Hank's Balanced Salt Solution (HBSS, Gibco) by centrifugation at 350g for 10 minutes. The pellet was resuspended in 3mL ACK lysis buffer (Quality Biological), followed by immediate addition of 7mL HBSS to dilute the leukocyte suspension. After centrifugation at 350g for 5 more minutes, the pellet was resuspended in HBSS containing $CaCl_2$ and $MgCl_2$ (Gibco).

Purified human neutrophils (2-3 x $10^7$) were plated on 10 mm petri dish for 30 minutes at 37°C, and then treated with phorbol 2-myristate 13-acetate (100 nM) for 4 hours at 37°C. Cells were washed with PBS, and DNase1 (4U/mL) was added for 20 minutes at 37°C to remove NETs from cell bodies. EDTA (5 mM final, ThermoFisher Scientific) was added to inactivate the DNase1. The supernatant containing NETs was centrifuged at 3000g for 10 minutes at 4°C to remove cell debris. The supernatant was collected and stored at -20°C. Nucleic acid concentration was measured using a Nanodrop™ spectrophotometer (ThermoFisher Scientific) as per manufacturer's instructions.

*Genomic DNA extraction from human neutrophils*

Purified human neutrophils from 50 mL whole blood were resuspended in HBSS prior to cell lysis with 10 mM Tris HCl (pH 8), 10 mM EDTA, 10 mM NaCl and 0.5% SDS, and incubated with proteinase K (10 mg/mL, Roche) at 37°C overnight as described[3, 4]. Equal volume of ultrapure phenol/chloroform/isoamyl alcohol (25:24:1, ThermoFisher Scientific) was added with vigorous swirling to mix for 15 minutes at RT, then centrifuged at 3000g for 10 minutes at 4°C. The aqueous supernatant was very slowly transferred to a new tube containing 2.5 X volume of 100% ethanol (Pharmco) to gentle intermix the layer by slow swirling to precipitate DNA. Strands of genomic DNA were removed from solution using a clean glass rod. The DNA was washed with 70% ethanol and centrifuged at 3000 x g for 10 min at RT. The DNA pellet was air dried and resuspended in Tris-EDTA buffer (pH 8) overnight at 4°C, then stored at -20°C. Nucleic acid concentration was measured using the Nanodrop™ spectrophotometer.

*Intravenous DNA administration into mice*

Wild-type (WT) C57BL/6 or littermates deficient in murine PF4 ($cxcl4^{-/-}$) mice were given a single bolus of HMW DNA (100 µg/mouse) via tail vein prior to blood collection from the retro-orbital sinus 30 minutes post DNA infusion into 1mM EDTA. Mice were euthanized 4 hours post DNA infusion, and blood was collected from the inferior vena cava into EDTA. Platelet-poor plasma was isolated by centrifugation at 2000g for 20 minutes at RT for study.

#### SUPPLEMENTAL FIGURES

**Figure S1. Size separation of HMW and digested NETs and DNA.**

Representative images of DNA size separation. (**A**) 0.4% agarose gel size-separation of intact 10 µg genomic DNA and NETs with M = 50 kb size marker. (**B**) NETs and DNA were digested with DNase1 and size-separated on a 0.9% agarose gel with M (left) = 1 kb size marker and M (right) = 100 bp size marker. (**C**) Same as in (B) but following digestion with the indicated restriction enzymes. Af2 = AflII, BsGrI-HF = BG1 and Al1 = AluI.

**Figure S2. Digested DNA-induced fibrin generation in PNP in the presence of PF4 and KKO.**

(**A**) Lag time for digested DNA-induced fibrin generation in the presence of increasing PF4 concentrations and 10 µg/mL KKO. (**B**) Lag time for digested DNA-induced fibrin generation in the presence of low dose PF4 (1 µg/mL) and the indicated KKO concentrations. Data are mean ± SEM of 3 independent experiments with the N value shown in each bar.

**Figure S3. DNA-induced thrombin generation in normal plasma, and fibrin formation in depleted plasma.**

(**A**) Representative kinetic curves of thrombin generation over time, initiated by digested NETs (left) and DNA (right) in the absence or presence of PF4 and KKO. **(B**) Representative kinetic curves of fibrin generation in PNP or FXI- or FXII-depleted plasma in the absence or presence of digested DNA with or without added PF4 (20 µg/mL).

**Figure S4. Effects of PF4 in DNA-induced fibrin generation in depleted plasma supplemented with FXI and FXII.**

(**A**) Lag time of digested DNA-induced fibrin generation and effects of PF4 in FXI-dep plasma supplemented with indicated FXI concentrations. (**B**) Lag time of digested DNA-induced fibrin generation and effects of PF4 in FXII-dep plasma supplemented with various FXII concentrations. Data are mean ± SEM of 3 independent experiments with the N value shown in each bar.

**Figure S5. PF4 protects endothelial cells from LMW dsDNA and ssDNA-induced VWF release mediated by TLR9.**

(**A** and **B**) Representative images of HUVECs showing immunofluorescent staining for VWF (red) and nuclei (blue) upon exposure to dsDNA (**A**) or ssDNA (**B**) fragments in the absence or presence of PF4. Mean fluorescence intensity (MFI) of released VWF from HUVECs exposed to dsDNA (**C**) or ssDNA (**D**), with or without the TLR9 inhibitor HCQ. Data were normalized to MFI of untreated cells without exposure to dsDNA or inhibitor (as indicated by dotted lines). Exposure to α-thrombin (C) or TNFα (D) serves as a positive control. Data are mean ± SEM of at least 3 independent experiments with the N value shown in each bar.

#### Figure S1

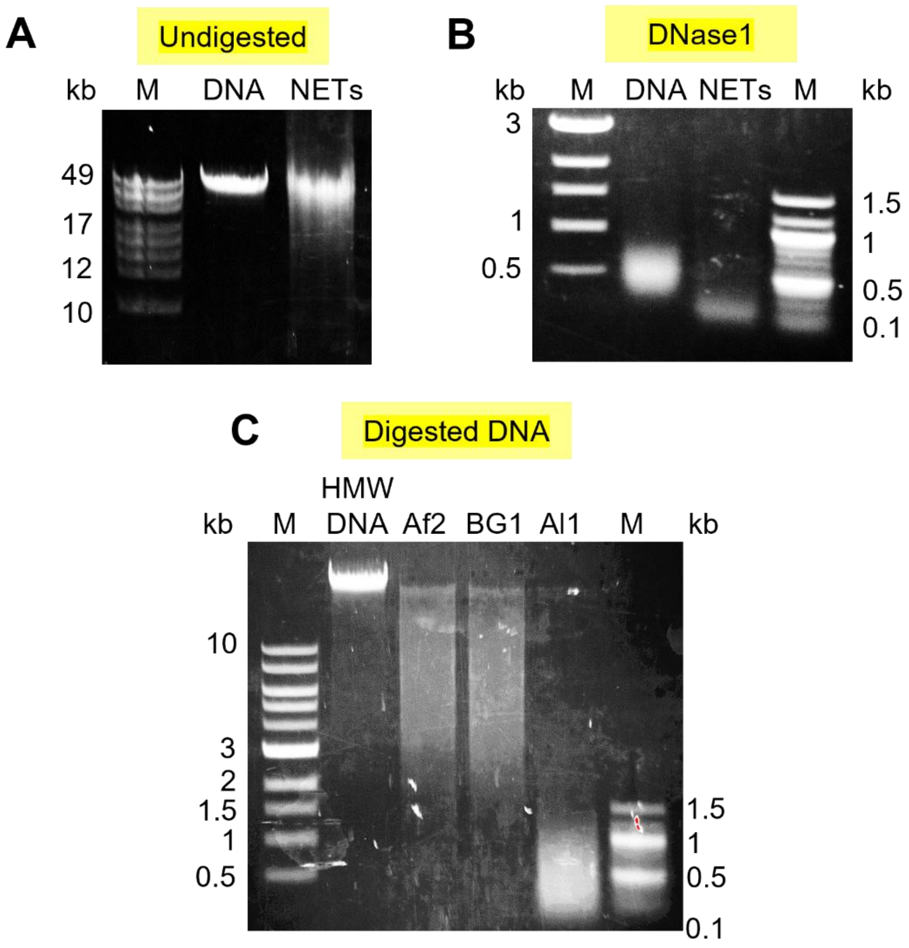
A
Undigested
kb
M
DNA
NETs
49
17
12
10
B
DNase1
kb
M
DNA
NETs
M
kb
3
1
0.5
1.5
1
0.5
0.1
C
Digested DNA
HMW
kb
M
DNA
Af2
BG1
Al1
M
kb
10
3
2
1.5
1
0.5
1.5
1
0.5
0.1

Figure S2

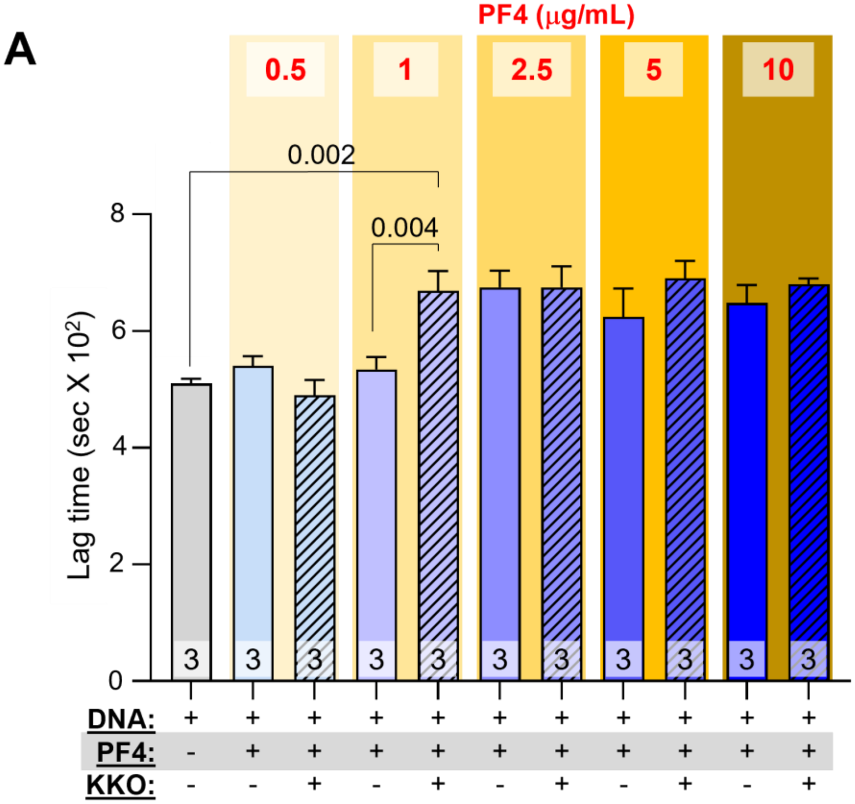
A
PF4 (μg/mL)
0.5
1
2.5
5
10
0.002
0.004
Lag time (sec X 10²)
8
6
4
2
0
3
DNA:
PF4:
KKO:

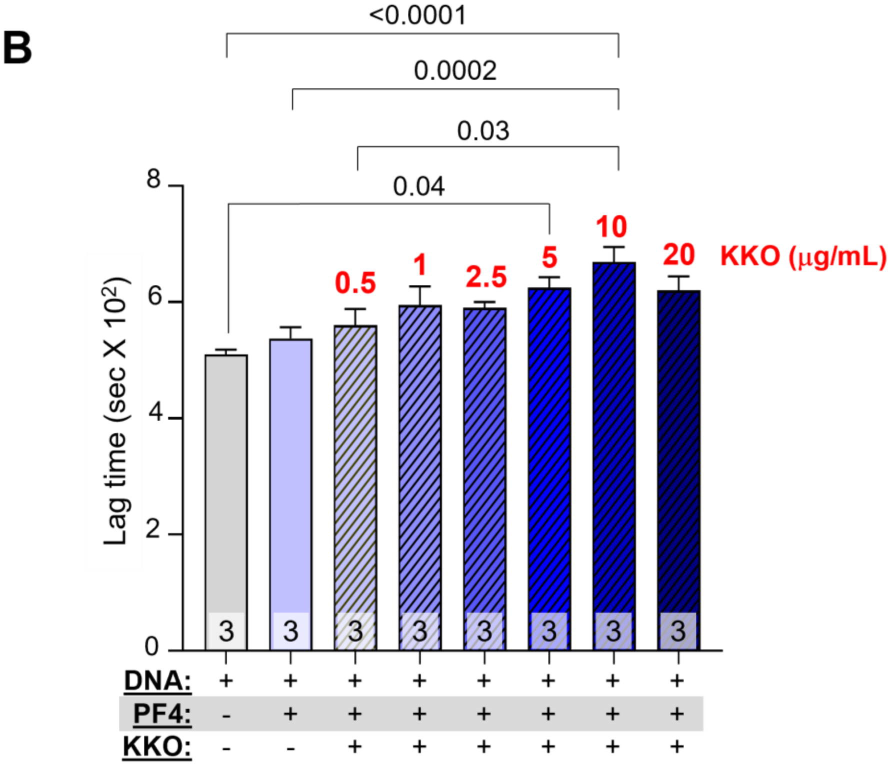
B
<0.0001
0.0002
0.03
0.04
Lag time (sec X 10²)
0.5
1
2.5
5
10
20
KKO (μg/mL)
8
6
4
2
0
3
DNA:
PF4:
KKO:

Figure S3

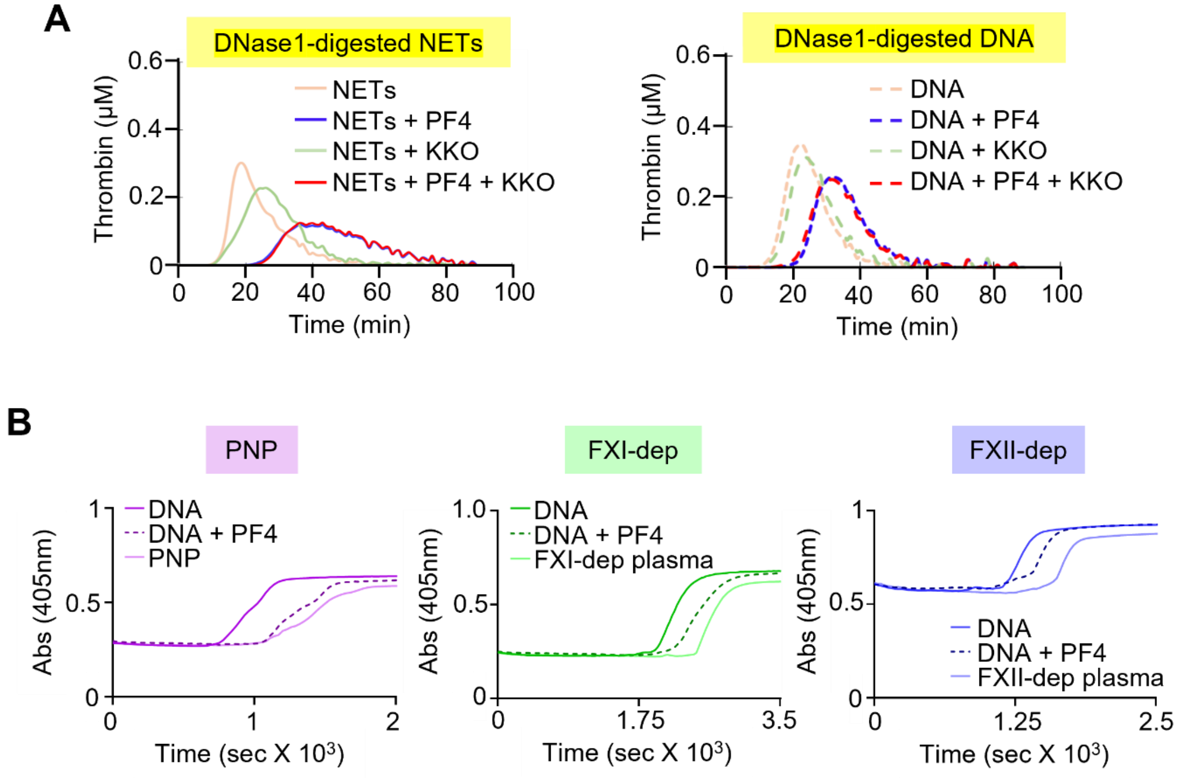
A
DNase1-digested NETs
Thrombin (µM)
Time (min)
NETs
NETs + PF4
NETs + KKO
NETs + PF4 + KKO
DNase1-digested DNA
DNA
DNA + PF4
DNA + KKO
DNA + PF4 + KKO
B
PNP
FXI-dep
FXII-dep
Abs (405nm)
Time (sec X 10³)
DNA
DNA + PF4
PNP
FXI-dep plasma
FXII-dep plasma

Figure S4

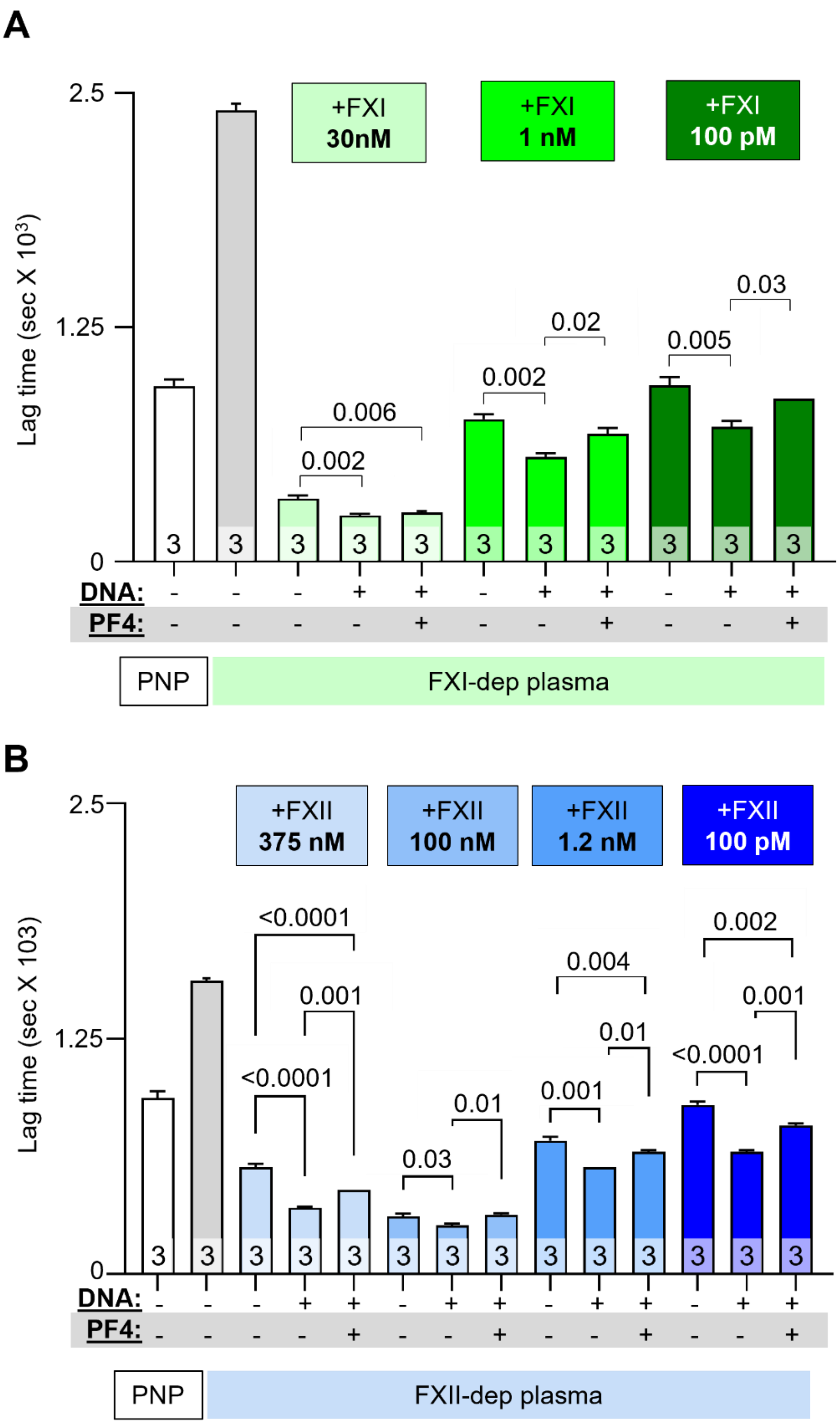
A
2.5
1.25
0
Lag time (sec X 10³)
+FXI 30nM
+FXI 1 nM
+FXI 100 pM
0.006
0.002
0.002
0.02
0.005
0.03
3
DNA: - - - + + - + + - + +
PF4: - - - - + - - + - - +
PNP
FXI-dep plasma
B
2.5
1.25
0
Lag time (sec X 103)
+FXII 375 nM
+FXII 100 nM
+FXII 1.2 nM
+FXII 100 pM
<0.0001
0.001
<0.0001
0.03
0.01
0.004
0.01
0.001
0.002
0.001
<0.0001
3
DNA: - - - + + - + + - + + - + +
PF4: - - - - + - - + - - + - - +
PNP
FXII-dep plasma

### Figure S5

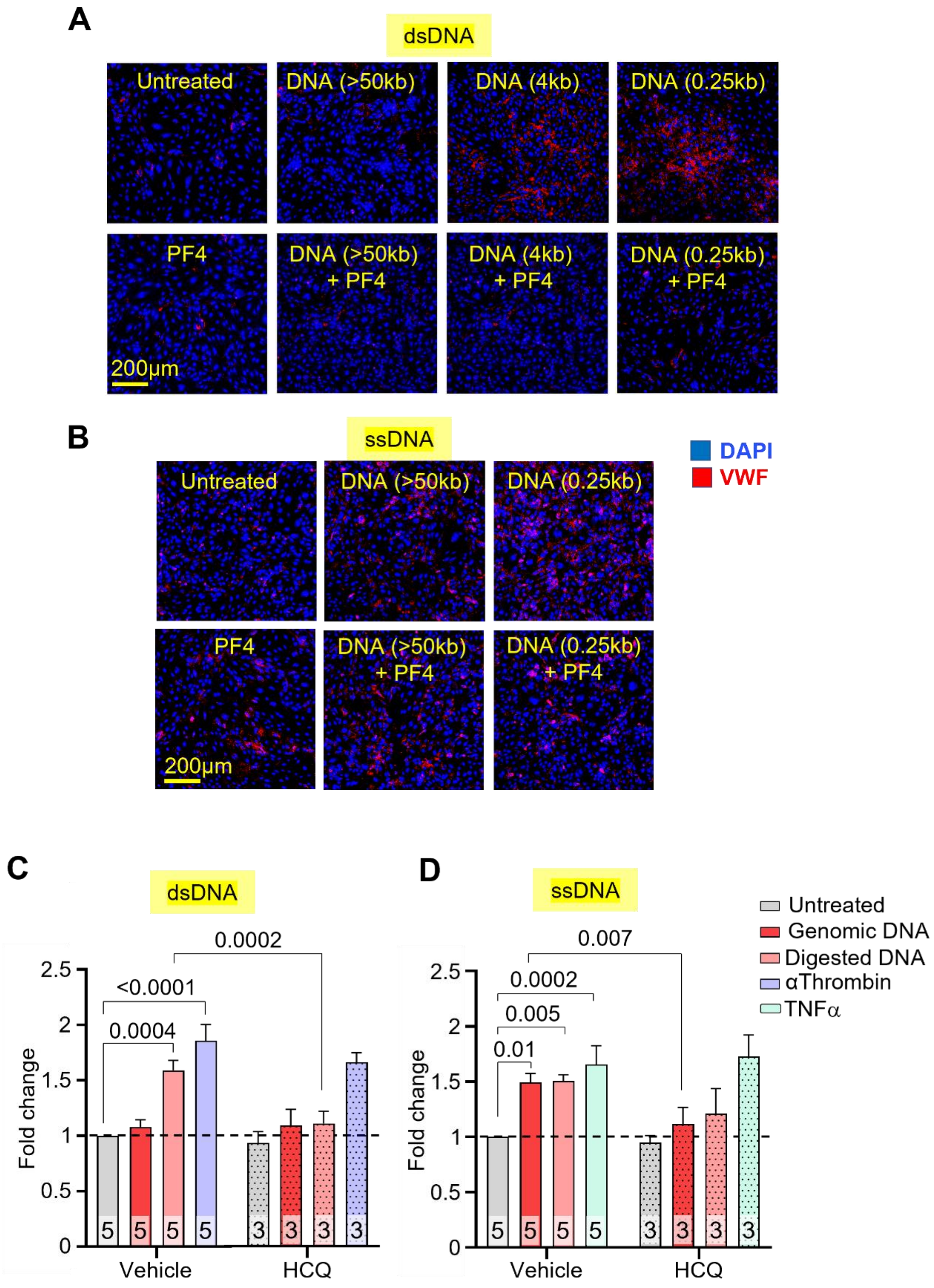
A
dsDNA
Untreated
DNA (>50kb)
DNA (4kb)
DNA (0.25kb)
PF4
DNA (>50kb) + PF4
DNA (4kb) + PF4
DNA (0.25kb) + PF4
200μm
B
ssDNA
DAPI
VWF
Untreated
DNA (>50kb)
DNA (0.25kb)
PF4
DNA (>50kb) + PF4
DNA (0.25kb) + PF4
200μm
C
dsDNA
0.0002
<0.0001
0.0004
Fold change
Vehicle
HCQ
D
ssDNA
0.007
0.0002
0.005
0.01
Fold change
Vehicle
HCQ
Untreated
Genomic DNA
Digested DNA
αThrombin
TNFα
